# Quantitative Characterization of Biological Age and Frailty Based on Locomotor Activity Records

**DOI:** 10.1101/186569

**Authors:** Timothy V. Pyrkov, Evgeny Getmantsev, Boris Zhurov, Konstantin Avchaciov, Mikhail Pyatnitskiy, Leonid Menshikov, Kristina Khodova, Andrei V. Gudkov, Peter O. Fedichev

## Abstract

We performed a systematic evaluation of the relationships between locomotor activity and signatures of frailty, morbidity, and mortality risks using physical activity records from the 2003 – 2006 National Health and Nutrition Examination Survey (NHANES) and UK BioBank (UKB). We proposed a statistical description of the locomotor activity tracks and transformed the provided time series into vectors representing physiological states for each participant. The Principal Components Analysis of the transformed data revealed a winding trajectory with distinct segments corresponding to subsequent human development stages. The extended linear phase starts from 35 − 40 years old and is associated with the exponential increase of mortality risks according to the Gompertz mortality law. We characterized the distance traveled along the aging trajectory as a natural measure of biological age and demonstrated its significant association with frailty and hazardous lifestyles, along with the remaining lifespan and healthspan of an individual. The biological age explained most of the variance of the log-hazard ratio that was obtained by fitting directly to mortality and the incidence of chronic diseases. Our findings highlight the intimate relationship between the supervised and unsupervised signatures of the biological age and frailty, a consequence of the low intrinsic dimensionality of the aging dynamics.

## INTRODUCTION

An accurate and non-invasive quantification of the aging process is essential for successfully translating basic research in the field of aging into future clinical practice. Most studies of aging in model organisms involve direct measurements of lifespan to characterize pro-or anti-aging effects of gene variants, nutrition conditions, or experimental therapies. In longer-lived animals, such as mammals, and especially in humans, the analysis of longevity itself is generally impractical since it would require long experiments with prohibitively large cohorts. Aging is a continuous phenotypic change and, therefore, one may alternatively hope to relate the dynamics of physiological variables representing the state of the aging organism to the incidence of diseases, frailty, and lifespan. Many markers of aging are shared between mice and humans and hence can be used to build a universal frailty index as a tool to quantify aging in preclinical studies [1–3]. Other useful metrics of aging include health span, maximum lifespan, and biological age [4]. The latter is commonly trained to predict chronological age from physiological measurements. These linear predictors, however, often fail to fully capture signatures of mortality and the incidence of diseases. This deficiency can be addressed with the help of log-linear mortality risk predictors, which have been used as proxies to quantify aging progress [5, 6]. It remains to be seen, however, if and how any of these measures of aging are related to each other in human populations, and whether the same associations hold true and therefore can be reliably examined in animal models.

The recent explosion in popularity of web-connected wearable devices has generated massive amounts of high quality measurements, including physical activity tracks, heart rate, skin temperature, etc., and consequently has created an unparalleled opportunity for aging research. It is projected that 400. such devices will be in use worldwide by 2020 [7] producing a deluge of biological data collected over many years. In this study, we performed a systematic evaluation of the relationship between locomotor activity and biological age, mortality risk, and frailty using human physical activity records from the 2003 – 2006 National Health and Nutrition Examination Survey (NHANES) and UK BioBank (UKB) databases. These large databases contain uniformly collected digital activity records provided by wearable monitors as well as health and lifestyle information, and death registry. We proposed a statistical description of 7-day long locomotor activity tracks and performed a Principal Components Analysis (PCA) of the study participants physical activity. This revealed that human life history is a continuous process. The explicit turning points on the aging trajectory signify marked changes in the character of the physiological state dynamics with age and correspond to the boundaries between the human development and aging phases. According to the Gompertz law [8], the mortality rate in human populations increases exponentially starting at forty years old. Therefore, we identify the distance travelled along the aging trajectory by an individual since the age of forty years old as the natural definition of biological age, or bioage.

The bioage variable describes most of the variance of the physical activity state and increases approximately linearly as a function of age. We found that biological age acceleration, i.e. the difference between the bioage of an individual and the corresponding age-and gender-matched cohort mean, is significantly associated with frailty and is also predictive of the remaining healthspan (the latter defined as the age of onset of prevalent chronic age-related diseases, such as coronary heart disease, including angina pectoris and heart attack, heart failure, stroke, hypertension, diabetes, and cancer) and lifespan. In the healthy individuals, therefore, the bioage acceleration is associated with activities that modify the lifespan, e.g. smoking, in such a way that smoking cessation leads to a reversible reduction of the bioage acceleration. A direct comparison shows that the unsupervised version of the biological age from the PCA correlates well with the negative logarithm of the averaged daily activity and with a log-linear proportional hazard predictor, trained to estimate mortality or morbidity from the same data. Finally, we investigate and highlight the intimate relationship between the supervised and the unsupervised biological age acceleration obtained from physiological variables on one hand, and traditional frailty assessment techniques on the other hand, as a direct consequence of the the high degree of correlation and hence the redundancy of physiological variables.

## RESULTS

### Quantification of Human Locomotor Activity

For this study, we used two large-scale repositories of wearable accelerometer track records made available by the 2003 – 2006 National Health and Nutrition Examination Survey (NHANES, 12053 subjects, age range 5 – 85 years old) and the UK Biobank (UKB, 95609 subjects, age range 45 – 75 years old). For both NHANES and UKB, a continuous, 7-day long activity track was collected for each subject, as well as data for a comprehensive set of clinical variables and death records up to nine years following the activity monitoring. Human physical activity is usually collected in the form of a series of direct sensor readouts, such a 3D accelerations, sampled at a specified frequency of time. However, the NHANES database provides sequences of transformed variables such as the number of steps and the activity counts per minute. Fig. 1A shows plots of two representative 2-day long activity tracks from a middle age (age 43) individual and an older (age 65) individual, who displayed the same level of overall activity. However, their patterns of activity were qualitatively different; the transitions between the different levels of physical activity appeared to be random. Therefore, instead of trying to determine the precise shape of the activity time series, we chose to apply a Markov chain approximation, which is a simple yet powerful probabilistic model from stochastic processes theory (see [9] for a review of its applications, including the stochastic modeling of biological systems).

**Fig. 1:**
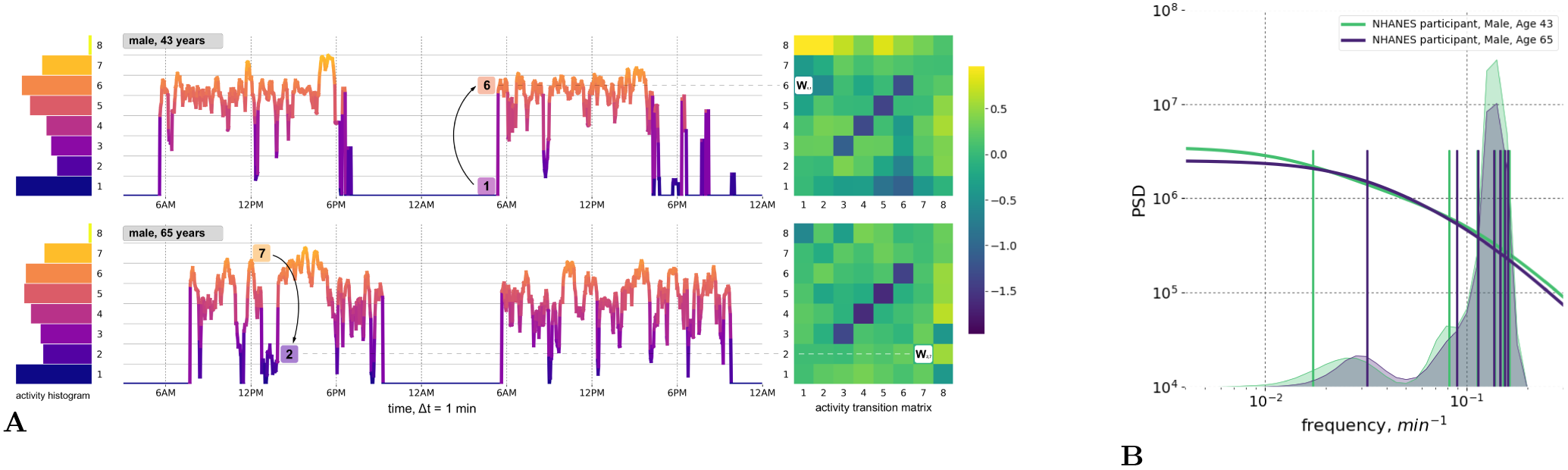
Quantitative Description of Human Locomotor Activity Tracks. **A.** Individuals with the same daily average level of activity can yet differ by their chronological age, health status and activity distribution during the day. Representative 2-day long locomotor activity tracks of two NHANES 2003 2006 cohort participants aged 43 (upper) and 65 (lower) illustrate how movement patterns can be visually different while having the same level of daily average activity. We quantify individual sample by dividing activity levels into 8 bins (left panel, histograms) and then counting the probabilities *W_ij_* of random jumps from each discrete activity state *j* to every other state. per unit time (right panel, color corresponds to intensity of transitions with respect to the population average). **B.** The eigenfrequencies of the Markov chain transition matrices are calculated for same two middle-aged and old individuals and represented by vertical bars (note the difference in the positions of the bars). The distribution of the eigenfrequencies in the relevant age-cohorts of 35 – 45 y.o. 65 – 75 y.o. are illustrated by overlaid transparent histograms (the light green and dark blue, respectively). Power Spectral Densities (PSD) reconstructed for Markov chain transition matrices (see Appendix A for details) reproduces the approximately a scale-invariant segment of the true PSD of the signal on time-scales up to tens of minutes. This characteristic shift of the cross-over frequency with age has been reported in numerous studies of human and animal locomotor activity (see text).

A statistical description of the participants’ activity was based on the concept that any future state of a Markov chain is determined only by its current state and the probabilities of transitioning between different states. We discretized the physical activity measurements over time into eight bins representing activity states (numbered from 1 to 8 and corresponding to increasing activity levels; see histograms to the left of the activity tracks in Fig. 1A). We counted the transitions between every consecutive pair of activity states along the track. For every pair of states *i* and *j*, the number of transitions from state *j* to *i* was then normalized to the number of times that state *j* was encountered along the entire activity record. This calculation yielded the kinetic transition rate, i.e. the probability of a stochastic “jump” from state *j* to state *i* per unit time. We then combined these transition rates into the transition matrix (TM) elements (shown as bins in heatmaps to the right of the activity tracks in Fig. 1A). The TM elements represent a complete description of the underlying Markov chain model (see Materials and Methods and the Fig. 1A description for additional details).

On a more technical level, the TM element values have the meaning of transition rates and hence can be related to the time scales characterizing the organism’s responses to external perturbations. To make this connection, we checked explicitly that the TM elements satisfied a detailed balance condition [10] and hence the TM eigenvalues represent inverse equilibration times. Using the relation between the autocorrelation function of the time series and the Markov chain TM from Appendix A, we plotted a reconstructed a Power Spectrum Density (PSD) in Fig. 1B for the physical activity signals corresponding to the same two study participants shown in Fig. 1A. Fig. 1B also shows the discrete sets of TM eigenvalues (the TM spectra) for the same individuals. The cross-over frequency on the PSD plots coincides with the lowest TM eigenvalues, corresponding to a time scale in the range of tens of minutes. The time scale corresponding to these eigenvalues is considerably longer than any period associated with body motion and, therefore, should reflect the organism’s physiological state. The observed decrease of the limiting time scale with age (see the density of the TM eigenfrequencies distributions for cohorts of 35 – 45 y.o. and 65 – 75 y.o. individuals in Fig. 1B) signifies a reduction of temporal correlations of physical activity in older subjects.

A transition matrix is a conceptually simple and intuitive aggregate characteristics of physical activity time series. TM elements are kinetic transition rates and the spectral properties of TM are directly related to the organism’s responses at physiological time scales. Therefore, TM elements calculated for each sample are a set of useful descriptors characterizing the physiological state of an organism and will be referred to as the physiological variables or, collectively, a physiological state vector or the organism state representation.

### Human Locomotor Activity Reveals Aging Trajectory

To reveal the intrinsic structure of the physical activity data for the entire NHANES study population, we used Principal Components Analysis (PCA), which is commonly used for multivariate data analysis and visualization [11]). PCA is an unsupervised method and can be employed without prior assumptions regarding the functional dependence of the biologically relevant variables on age. Fig. 2A shows the distribution of the transformed data along the first three PCs, *PC*_1_ vs. *PC*_2_ vs. *PC*_3_. Each point represents the average activity profile representing the age-matched cohorts of men and women. The physiological state vector changes in the course of human lifespan, meaning the aging trajectory is continuous, and can be subdivided into distinct phases that are recognizable as the subsequent human development phases. We used one of commonly accepted systems of age classification [12] and divided the trajectory into four segments: (I) childhood and adolescence (younger than 16 years old); (II) early adulthood (16 – 35 y.o.); (III) middle ages (35 − 65 y.o.); and (IV) older ages (older than 65 y.o.). Although there was a significant difference between the trajectories of male and female participants, their overall shape and direction were relatively similar.

**Fig. 2:**
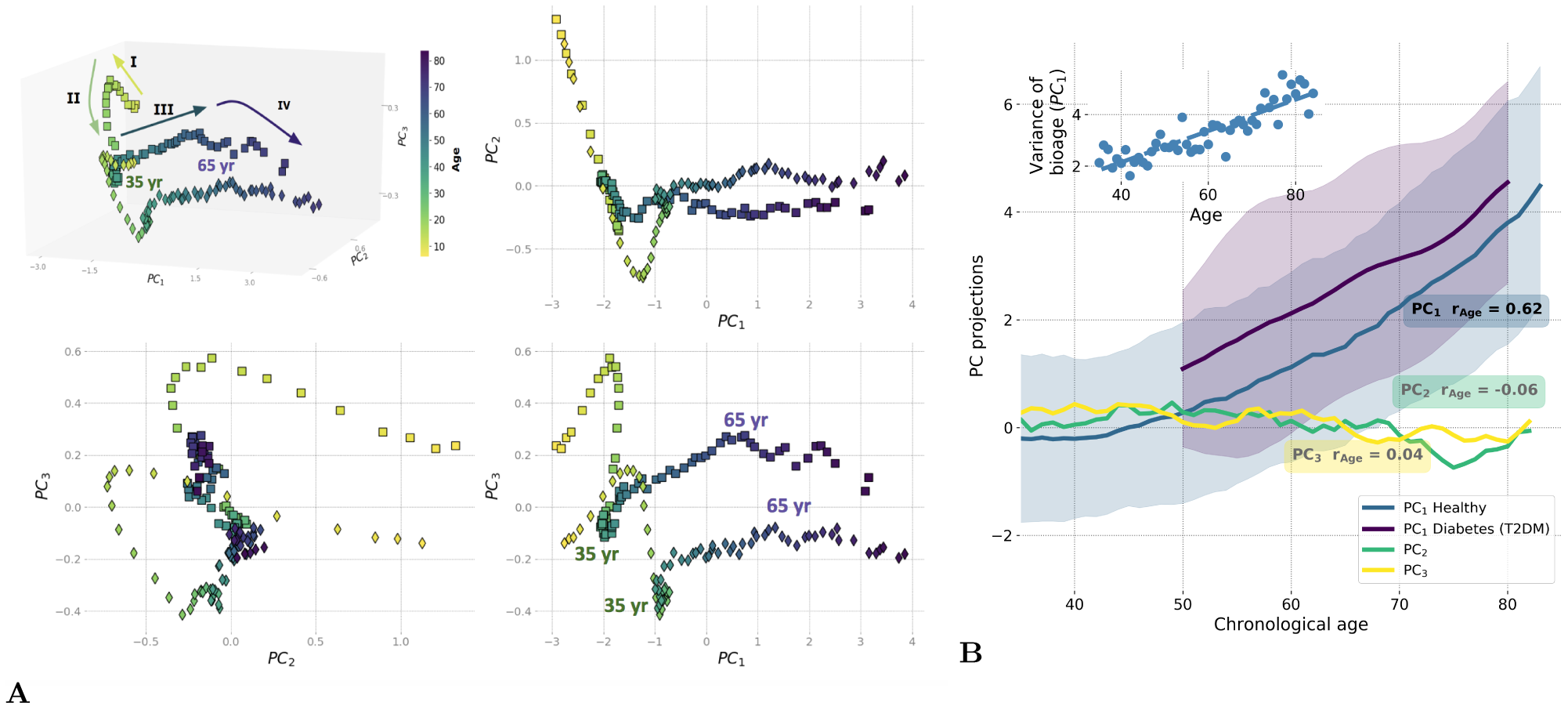
Principle Component Analysis (PCA) reveals low-dimensional aging trajectory. **A.** The graphical representation of the PCA for 5 – 85-year-old NHANES 2003 – 2006 participants follows a winding aging trajectory. Samples were plotted in the first three PCs in 3D space along with 2D projections. To simplify the visualization, the PC scores are shown for the age-matched averages for men (squares) and women (diamonds) and color-coded by age. The Roman numerals and corresponding arrows illustrate the approximately linear dynamics of PC scores over sequential stages of human life: I) age*<* 16; II) age 16 − 35; III) age 35 − 65; and IV) age *>* 65. **B.** Age-dependence of PCA scores along chronological age for NHANES 2003 − 2006 cohort aged 35+ is shown by age-cohort average values. Human physiological state dynamics has a low intrinsic dimensionality: only the principal component score, *PC*_1_, which corresponds to the largest variance in data, showed a notable correlation with age (Pearson’s *r* = 0.62 for *PC*_1_ and *r <* 0.2 for other PCs) and therefore could be used as a natural biomarker of age. Shaded regions illustrate the spread corresponding to one standard deviation in each age-matched cohort for *PC*_1_. The inset shows the increase of variance in biological age (*PC*_1_) in the age-and sex-matched cohorts along the chronological age.

According to the Gompertz law, the risk of mortality in human populations increases exponentially in mid-life, starting around age 40 [8, 13]. We observed the relevant turning point, a significant shift in character of the physiological state dynamics, between the aging trajectory of segments II and III exactly at this age (Fig. 2A), corresponding to the transition from early adulthood to middle age. In addition, we found another cross-over at approximately 65 years old, corresponding to the boundary between middle age and older age (segments III and IV in Fig. 2A), and occuring in the vicinity of the average human healthspan, defined as the survival free of chronic age-related diseases. According to a recent World Health Organization report [14], the average healthspan is approximately 63 years old. Since aging is the focus of this study, we limited the subsequent analysis to participants older than 40 years old.

In this restricted dataset, aging manifested itself as the approximately linear evolution of the participants’ physiological state along the *PC*_1_ direction, which by definition is the direction of the most variance in the data. Only *PC*_1_ scores in this group of participants were strongly associated with chronological age (Pearson’s correlation coefficient *r* = 0.62; see Fig. 2B). Based on our observations, we propose that the first PC score, PC_1_, represents a natural definition of biological age, or bioage, which is a quantitative measure of the aging process in the most relevant age range. It increases linearly with chronological age for participants older than 40, as shown in Fig. 2B. In addition, its correlation with age persists (*r* = 0.47) even in the cohort of the most healthy individuals (according to an implementation of the Frailty Index adopted for NHANES in [2]), suggesting that the association cannot only be attributed to the development of illness. The non-linearity in the bioage dynamics with age is weak and is not sufficient to explain the exponential growth of mortality risks with age. The exponential fit of our data yields the doubling rate of approximately 0.02 per year, which is far less than the doubling rate of 0.085 per year according to the Gompertz law. Hereafter, we refer to the biological age dynamics (the dependence of the biological age on the chronological age) and the associated variation of the physiological variables with age as “aging drift”.

To characterize the effect of diseases on the dynamics of biological age, we assessed the effect of type 2 diabetes mellitus (T2DM), a common age-related disease, on biological age as defined by *PC*_1_. We compared the mean and standard deviation of biological age in age-matched cohorts of T2DM patients and healthy subjects (Fig. 2B). Generally, the T2DM patients appeared to be older according to their biological age (*PC*_1_) when compared to their chronological age-matched peers. The biological age difference between the healthy and T2DM groups did not significantly change with chronological age.

### Biological age acceleration predicts mortality and the incidence of chronic diseases (morbidity)

The biological age acceleration (BAA) is commonly defined as the difference between the biological and the chronological age of an individual. It is a natural measurement of a person’s aging process relative to that of their peers, and can be associated with their lifespan and the presence of chronic age-related diseases (see [15–17]). We propose a more general and robust definition of aging acceleration associated with an arbitrary variable: namely, the residual from the average of the same variable in a cohort of age-and gender-matched individuals (see a recent example in [5]). In this way, aging acceleration can be calculated for any measurement, not simply those expressed in years of life, but also for more sophisticated measures of aging progress, including “biological age” *PC*_1_.

First, we tested the hypothesis that the BAA of “biological age” *PC*_1_ is associated with all-cause mortality. We used the death records available for 4612 NHANES participants aged 40+, including 550 observed death events during the followup of up to 9 years, and obtained the BAA by adjusting the *PC*_1_ score for gender and chronological age. A Cox proportional hazard regression using the BAA value as a co-variate yielded the hazard ratio HR = 1.58. 95% CI = 1.54−1.62, see Table I. Notably, we observed a prominent correlation between *PC*_1_ and the average level of physical activity, specifically its log-scaled value (Fig. 3B). As expected, the BAA of the measurement of total activity was also significantly associated with mortality in the same dataset.

**Fig. 3:**
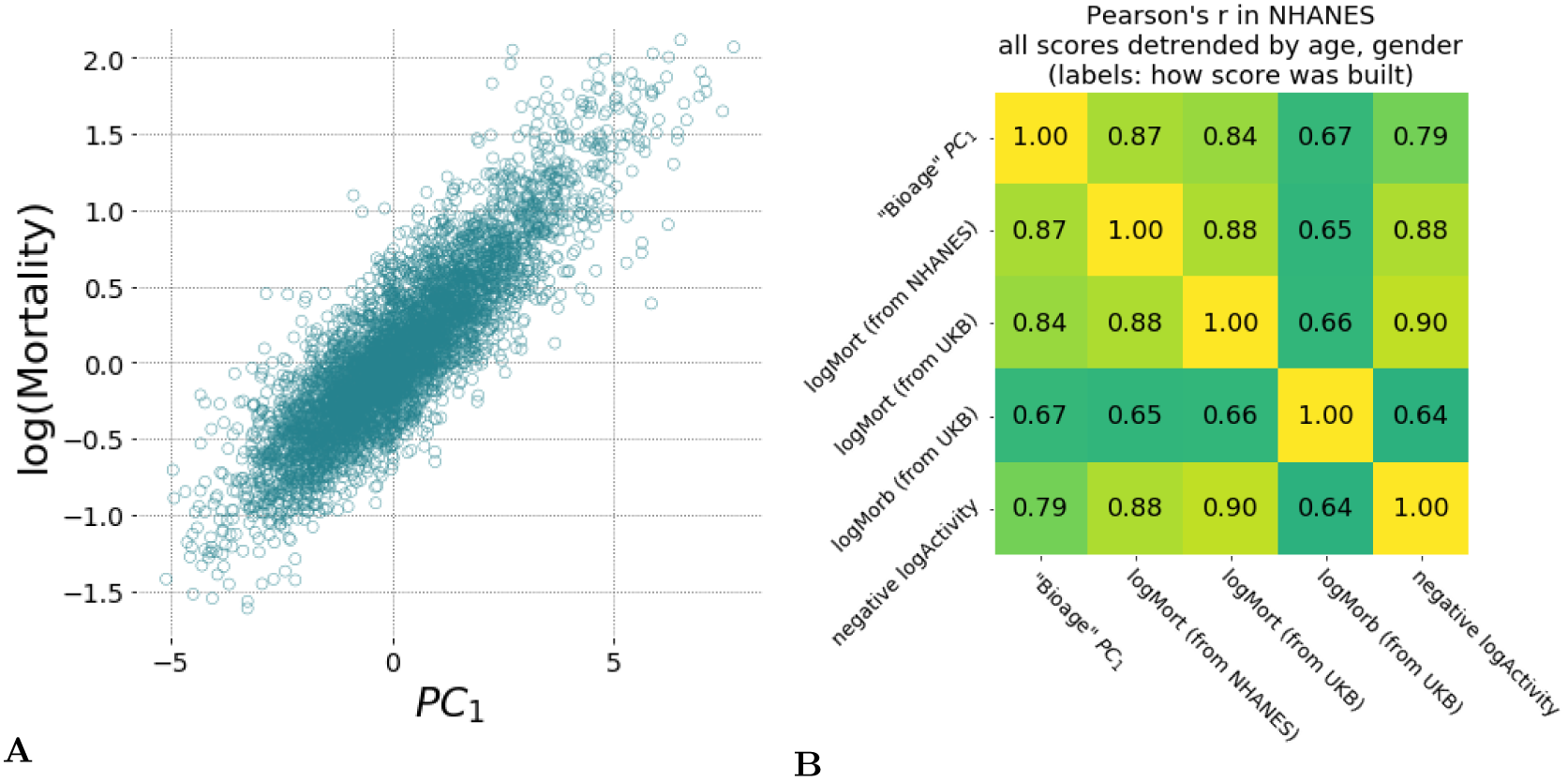
Hazards ratio models show high correlation with each other and are strongly associated with average level of physical activity and the largest variance in physiological measurements (*PC*_1_). **A** Scatter plots of estimated mortality hazard ratio (log-scale) vs *PC*_1_ scores and log-hazard ratio estimated by “LogMort” model trained on NHANES survival follow-up data shows high correlation (Pearson’s *r* = 0.87). **B.** Different models for hazard ratio of mortality and morbidity show high correlation between each other and the *PC*_1_ “biological age” in NHANES samples. Models for mortality and morbidity were built using Cox proportional hazards method based on either NHANES or UKB death follow-up data and UKB follow-up on diagnosis. All values were adjusted by age and gender and thus represent the corresponding Biological Age Acceleration (BAA) values.

**TABLE I:** Association of the biological age *PC*_1_ and the log-hazard ratio mortality risk estimation with prospective mortality and morbidity events calculated using Cox-proportional hazard models with adjustment for age and gender.

| Tested model | Dataset (kind of prospective events) | HR (95% CI) | p-value |
| --- | --- | --- | --- |
| "Bioage" $PC_1$<br>(adjusted for age, gender) | NHANES (mortality) | 1.57 (1.53 - 1.61) | $p < 1E - 10$ |
| | UKB (mortality) | 1.81 (1.76 - 1.86) | $p < 1E - 10$ |
| | UKB (morbidity) | 1.16 (1.13 - 1.19) | $p = 2.4E - 7$ |
| "LogMort": log-hazard ratio<br>(adjusted for age, gender) | NHANES (mortality) | 1.76 (1.73 - 1.79) | $p < 1E - 10$ |
| | UKB (mortality) | 1.81 (1.76 - 1.86) | $p < 1E - 10$ |
| | UKB (morbidity) | 1.15 (1.12 - 1.18) | $p = 1.7E - 6$ |

We confirmed the association of “biological age” *PC*_1_ with mortality risks using the independent UKB dataset, which encompasses another 93597 individuals aged 43 − 78 years (mean of 62.4 ± 7.8 yr) with 285 recorded deaths during a 3 year follow-up. We estimated the “biological age” *PC*_1_ scores for UKB participants using the same Principal Component vector obtained in the NHANES dataset and found a highly significant contribution in the Cox proportional model HR = 1.81. 95% CI = 1.76 − 1.86. This suggests that the biological age signature is relatively robust and can be applied to different datasets.

BAA also turned out to be a significant risk factor for the prospective incidence of chronic related diseases. Following [18], we observed that there is a large cluster of age-related diseases, such as cardiovascular diseases (coronary heart disease, heart attack, heart failure, stroke), diabetes, hypertension, and cancer. All the diseases from the list are characterized by an exponentially increasing probability of incidence, with a doubling time similar to that of the mortality rate of eight years identified in the Gompertz mortality law in human population. Assuming the single underlying risk factor, *i.e.* the aging itself, we defined the healthspan as the age marked by the first diagnosis of any of the aforementioned diseases. To test the association between the BAA and the healthspan, we selected 43533 UKB participants without any age-related disease at the moment of locomotor activity assessment (1331 disease events during 3 year follow-up) and tested the association with the prospective first incidence of chronic disease using the Cox model. The observed hazard ratio (HR = 1.16. 95% CI = 1.13 − 1.19) demonstrated a highly significant effect (*p* = 2.4. 7; see Table I). These data suggest that BAA is a risk factor that marks the increased probability of the prospective incidence of age-related diseases in the healthy individuals.

### Supervised proportional hazards models and the biological age

The prevalence of mortality and/or the incidence of major diseases in a population can be inferred from the death records or clinical data and represent the ultimate objective measure of an individual’s resilience to disease. In this section, we introduce and characterize another natural biological age measure–the log-hazard ratio of a risk model–fitted directly to the experimentally observed occurrence of death or the incidence of chronic diseases. We started by confirming that the empirical mortality in the NHANES dataset follows the well-established Gompertz law. To do this, we fit the age-at-death statistics using a parametric Cox-Gompertz model, which is a version of the Cox-proportional hazard model with an explicit Gomprtz assumption on the follow-up mortality [19]. We obtained a mortality rate doubling constant of 0.08 per year, which is close to the empirical value of Γ ≈ 0.085 per year. This constant corresponds to a mortality rate doubling time of 8 years [20] and an average life expectancy of 75 years, which is close to 79, the reported value for the United States population (see [21]).

Having established that the expected pattern of exponential mortality exists in the NHANES dataset, we used a Cox proportional hazards model [22], with gender and all of the physiological variables obtained from the locomotor activity as covariates. The mortality risks model was trained on data for NHANES participants aged 40 and older and was then used to estimate the logarithm-scaled risk of mortality for participants from both NHANES and UKB datasets. For simplicity, we will denote these predicted risks as “logMort” to distinguish them from the “biological age” and other models utilized in this study. The risk of death was found to increase exponentially as a function of biological age (Fig. 3A; the determination coefficient of the corresponding log-linear model *R*^2^ = 0.81). This further supports our conjecture that the *PC*_1_ score is a quantitative measure of the biological aging progress. These data suggest that the logarithm of the risk of death may serve as a viable but essentially equivalent approach to evaluate biological age.

The estimated log-hazard ratio robustly predicted the risk of mortality and chronic diseases across the sex-and age-adjusted NHANES and UKB cohorts (see Table I for the “logMort” model summary). For every calculation, we used the log-hazard ratio predictor as a covariate and adjusted for chronological age and gender. The resulting BAA estimates were strongly associated with mortality risks in NHANES population (HR = 1.76. 95% CI = 1.72−1.80). The observations were confirmed in the independent dataset of UKB participants, with a hazard ratio similar to that obtained for the “biological age” *PC*_1_ after adjusting for age and gender (HR = 1.81. 95% CI = 1.76 − 1.86). The log-hazard ratio of the mortality model was also significantly predictive of the prospective morbidity (*p* = 1.7. 6), with a similar result observed for the unsupervised “biological age” *PC*_1_ in Table I.

The apparent similarity between the ability of the supervised (“LogMort”) and the unsupervised (*PC*_1_) biological age models to predict the incidence of death and age-related diseases is a consequence of the high degree of correlation characteristic for biological data and hence the redundancy in the physiological variables measurements. The concordance between the biological age predictors also hints at another practical opportunity to train bioage models. In most cases, the death records are scarce even in very large studies, since the mortality rate is small and requires a very long participant followup to gather sufficient information on the lifespan of the population. The incidence of disease is much higher than the mortality, since the healthspan is shorter than lifespan, and hence any dataset with aging human subjects would contain a considerable fraction of people suffering from age-related diseases. Given this information, we could build a hazards model of the incidence of disease in the UKB dataset, “LogMorb (UKB)”, using the individuals who were considered healthy at the time of the physical activity assessment. As expected, all of the presented mortality and morbidity risks model’s predictors, including the mortality ratio built in UKB data, “LogMort (UKB)”, and the unsupervised biological age *PC*_1_, demonstrated remarkably high correlations across samples after adjustment for age and gender in both NHANES (Fig. 3B) and UKB (data not shown). Our findings indicate a substantial overlap between the signatures of the mortality and morbidity risks that can be used to define human lifespan and healthspan, respectively.

### Biological Age Acceleration and Frailty

Finally, we checked the association of BAA with the Frailty Index (FI), a classical measurement of general health using functional and clinical information rather than physiological characteristics. In general, FI is proportional to the overall number of health deficits or diseases and is significantly associated with mortality, see, e.g., [1, 2]. We had already observed that the BAA computed using biological age is higher in groups diagnosed with T2DM, a disease whose presence increases the FI of the affected individual (see Fig. 2B). We made the same observation using the supervised biological age models (the “LogMort” model), in which the predicted mortality log-hazard ratio was significantly higher in the sex-and age-adjusted cohorts in the NHANES dataset that constituted the “frail” and “most frail” individuals (Fig. 4C) stratified according to a version FI tailored for NHANES dataset [2].

**Fig. 4:**
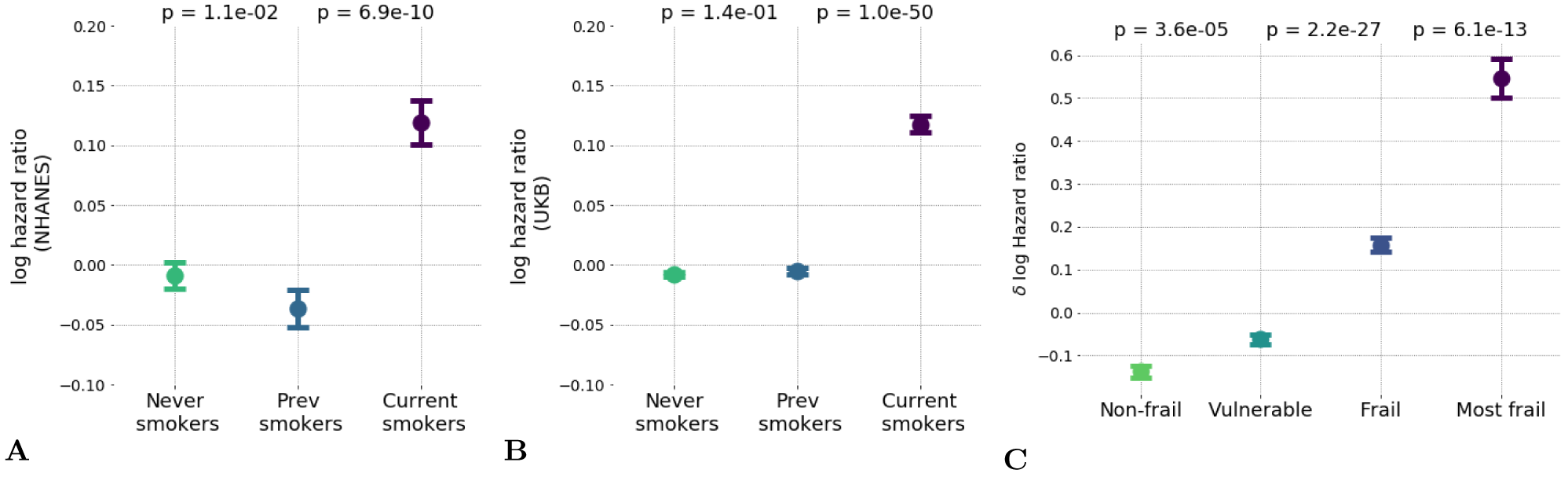
Hazards ratio model distinguished low and high-risk populations and hazardous lifestyles. A-B. The effect of unhealthy lifestyle such as smoking caused reversible effect on estimated hazards ratio in (A) the NHANES population and (B) the UK Biobank datasets; **C.** Distribution of logarithm of estimated hazards ratio in frailty cohorts shown by median ± standard error of mean (S.E.M.). “Frail” and “most frail” cohorts are stratified on the basis of the respective Frailty Index (FI) values computed according to [2] and are characterized by significant difference in the predicted log-mortality.

The BAA is a continuous measurement with a broader significance beyond predicting the degree of frailty. For the individuals from the healthiest cohorts (i.e. excluding individuals with the “frail” and “most frail” designations), the BAA was identified as a signature of having an elevated risk of chronic diseases incidence (see Table I) and death associated with a hazardous lifestyle. To demonstrate this association, we compared the biological age of current smokers, those who never smoked, and those who stopped smoking. The calculation revealed a significant difference of BAA between the individuals who currently smoke and those who never smoked (see Fig. 4A). Interestingly, the BAA level of those who stopped smoking was reversed to that of never-smokers. We validated this effect in the UKB population using the model trained only on the NHANES dataset, which produced the same arrangement of BAA differences (see Fig. 4B).

## DISCUSSION

In this study, we evaluated various indicators of age and frailty using a highly accessible measurement of human physiological state: the time series representing the accelerometer records of human physical activity. We used a number of multivariate data analysis techniques, including unsupervised Principal Component Analysis and supervised proportional mortality and morbidity hazard models, to evaluate distinct NHANES and UKB datasets. The phenotypic changes reflected by the aggregate physical activity variables used in this study and associated with the development and aging showed different dynamics depending on the life stages (Fig. 2A). We determined that the age range starting from 40 years and older, corresponds to the transition from early adulthood to middle age, and provided the most relevant information for an investigation into the dynamic origins of Gompertz mortality law in humans. The dynamics of the biomarkers of age can be described as a highly deterministic process. The aging trajectory can be identified with the help of PCA in a totally unsupervised way. We found that while the first PC score (*PC*_1_) increases significantly with age, the other PC scores are virtually independent of age. Therefore, *PC*_1_ has the meaning of the distance travelled along the aging trajectory and hence represents a natural measurement of biological age (Fig. 2B).

In the language of dynamical systems theory, the reduction of the physiological state dynamics to a continuous aging trajectory that was revealed by the PCA is a hallmark of the low intrinsic dimensionality of how physiological variables evolve with age. This situation is typical for the slowest biological processes, such as morphogenesis [23] and aging [24], that exhibit similar characteristics, including a critical period of slowing down, increased variance, and a strong correlation between key variables [25]. In [24], we suggested that the dynamics of the collective variable associated with the criticality is identifiable from the PCA and is the driving force behind the characteristic increase in mortality with age. The biological age associated with the first PC score is therefore an emergent organism-level property, the key indicator of the aging process. The aging at criticality hypothesis [24] is thus a reasonable theoretical explanation for the success and popularity of PCA for quantifying biological age in this study and in almost every other kind of biological signal (see, for example, [26–30]).

The biological age as defined by *PC*_1_ was found to increase linearly with chronological age in the NHANES dataset for individuals older than 40 as shown in Fig. 2B. The observed level of the non-linearity is weak and is insufficient to explain the exponential growth of mortality risks with age. The variance in biological age *PC*_1_ in age-and sex-matched cohorts in this population also increased linearly (see the inset in Fig. 2B). The latter could be a hallmark of diffusion, suggesting that the biological age variable not only drifts over time but also undergoes a random walk under the influence of stochastic forces. An alternative explanation would require a non-linear mode coupling between the aging drift and higher frequency modal variables characterizing fast responses of the organism state to random external and endogenous stress factors. Further experiments would be needed to confirm the veracity of these hypotheses.

The linear association of physiological variables with age is the reason behind the success of “biological clocks” that are commonly trained as linear predictors of chronological age, such as DNA methylation [15, 16], IgG glycosylation [31], blood biochemical parameters [17], gut microbiota composition [32], and the cerebrospinal fluid proteome [33]. To date, the “epigenetic clock,” based on the levels of DNA methylation (DNAm) [15, 16], appears to be the most accurate measurement of aging, showing a remarkably high correlation with chronological age. The DNAm clock predicts all-cause mortality in later life more accurately than chronological age [34]–and is elevated in groups of individuals with HIV [35], Down syndrome [36, 37], and obesity [38]–but is not correlated with smoking status [39].

The supervised linear predictors of chronological age that are built using physiological variables, including the DNA methylation clock, are trained to minimize the biological age acceleration (BAA), the difference between an individual’s predicted chronological age and their actual age. BAA itself is a sensible biological variable associated with mortality risk or disease status, and therefore refining the correlation to the chronological age often comes at the expense of losing biologically significant information. For example, some popular biological age models fail to fully capture signatures of all-cause mortality [5, 6]. This is consistent with the conclusions of a recent study where Frailty Index better predicted mortality rates compared to DNAm age [40]. Also, in a separate epigenome-wide association study, the reported DNAm signature of all-cause mortality was found to contain an extra component that was independent of the “epigenetic clock” [41].

An alternative approach to predicting chronological age involves directly estimating mortality risk from a set of physiological variables. Since mortality in human populations increases exponentially with age, the log-hazard ratio prediction is roughly a linear function of age and therefore represents a sensible supervised predictor (i.e. trained using the death registry information) of biological age [5, 6]. In the present work, we demonstrate that the BAA of the predictors of biological age using a log-hazard ratio correlates with the BAA of the principal component score. We therefore conclude that the both approaches yield highly concordant biological age estimations and, as such, represent the same underlying biology: both phenotypes are associated with Frailty Index.

In the most healthy (i.e. the least frail) individuals, the BAA turned out to be a signature of a response to generic stress and was associated with an elevated incidence of disease and mortality risk caused by hazardous behaviors such as smoking (see Figs. 4A and 4B). Our findings regarding the impact of smoking are in concordance with the earlier results obtained by [39], where the Frailty Index - but not a linear predictor of chronological age-significantly correlated with the regulation of smoking-associated methylation sites. We also showed that the BAA are significantly lower in individuals who had smoked early in life, but that the trend is reversed upon quitting smoking, presumably reducing the risks of future development of irreversible chronic health conditions. The seeming reversibility of the biological age variation associated with smoking aligns with the reported benefits of quitting smoking early in life on life expectancy [42]. This observation fuels the hypothesis that the effects of BAA can be modulated by lifestyle changes or therapeutic interventions.

We observed that the BAA was not only a significant risk factor of disease-associated mortality, as in [6], but also of the incidence of chronic age-related diseases in a prospective follow-up study. The latter was supported by a significant association between *PC*_1_ and the log-linear proportional incidence of chronic age-related diseases risk estimate in the UKB dataset. These findings corroborate the findings investigating the GWAS of healthspan [18], where at least some of the genetic variants associated with longer healthspans were found to predict both lifespan and the incidence of specific age-related diseases.

The overall level of physical activity decreases with age and predicts the extent of remaining lifespan in both humans and other species [43, 44]. We observed a high concordance between any of the biological age predictors and the negative logarithm of the average level of daily physical activity (Pearson’s *r* = 0.79, with the higher values of biological age corresponding to lower levels of physical activity). The average activity alone, however, cannot be a good single measurement of biological age or frailty, as the quantity depends strongly on lifestyle and hence may be poorly transferable across datasets, types of wearable devices, and diverse populations. This observation is illustrated by the distribution of average activity levels across countries [45], which does not correlate with life expectancy. For example, the average physical activity is approximately 50% higher in UKB participants compared to NHANES participants, and yet the average life expectancies are very similar.

To address these limitations, in this study, we turned to a richer set of physical activity characteristics, the TM components. These components provide a window into the autocorrelative properties of the physical activity time series, which enable the detection of repeated patterns, on physiologically relevant timescales (minutes to hours). More specifically, we found that the smallest TM eigenvalue increases with age, suggesting a gradual degradation of the long-time correlation of the movement patterns in older or frail individuals. This property is commonly observed in studies of aging and age-associated neurological and mental disorders both in humans [46] and animals [47], specifically in Alzheimer’s disease [48], depression [49], and bipolar disorder [50], and therefore could be attributed to increasing frailty irrespective of the corresponding disease.

We tested the robustness of the biological age models across the independent NHANES and UKB datasets, which differed not only in population (the United States vs. the United Kingdom) but also in the accelerometer sensor hardware. In these datasets, the aggregate characteristics that represented human physical activity demonstrated a remarkable degree of transferability. We used the models that were trained on the NHANES dataset to profile risks associated with lifestyles (such as smoking), the future incidence of age-related diseases, and the remaining healthspan of individuals from the UKB dataset. We found this observation reassuring and hope that the risk models can be further improved with the help of modern engineering; for example, convolutional neural networks are now capable of inferring longer correlations and can better capture non-linear relationships between the identified features of the signal (see [5]).

The robust identification of age and frailty biomarkers requires access to large-scale datasets that have been annotated with age, gender, historic and prospective clinical information and the death registry. Our work suggests that the variation among physiologically relevant variables is often the result of very few underlying factors, most notably frailty. We characterized a simple unsupervised (PCA-based) measure of the “biological age” in a novel signal derived from the physical activity track records from wearable devices. The model performed well using the minimum amount of information and can thus serve as a good initial estimate for a series of more sophisticated biomarkers of age and mortality risks. We hope that our work will bring necessary attention to electronic activity records and help demonstrate its potential for aging research and for broader health and wellness applications.

## CONCLUSION

In conclusion, we demonstrate a possibility to quantify time series of human physical activity. We show a possibility to extract locomotor activity-based signatures of life staging, aging acceleration, increased morbidity and mortality risk in association with diseases and hazardous lifestyles. We report the intimate relationship between the unsupervised measurement of biological age (the distance travelled along the aging trajectory), frailty, and the log-proportional hazard ratio of models trained to predict the risks of chronic diseases or all-cause mortality. On a more practical level, our findings highlight an opportunity for the deployment of fully automated wellness intelligence systems capable of processing tracker information and providing dynamic feedback in a completely ambient way. This could be used for improved engagement in health-promoting lifestyle modifications, disease interception, and clinical development of therapeutic interventions against the aging process.

## MATERIALS AND METHODS

### NHANES dataset

Locomotor activity records and questionnaire/laboratory data from the National Health and Nutrition Examination Survey (NHANES) 2003-2004 and 2005-2006 cohorts were downloaded from [www.cdc.gov/nchs/nhanes/index.htm]. NHANES provides locomotor activity in the form of 7-day long continuous tracks of “activity counts” sampled at 1*min^−^*^1^ frequency and recorded by a physical activity monitor (ActiGraph AM-7164 single-axis piezo-electric accelerometer) worn on the hip. Of 14,631 study participants (7176 in the 2003-2004 cohort and 7455 in the 2005-2006 cohort), we filtered out samples with abnormally low (average activity count *<*50) or high (*>*5000) physical activity. We also excluded participants aged 85 and older since the NHANES age data field is top coded at 85 years of age and we desired precise age information for our study. The mortality data for NHANES participants is obtained from the National Center for Health Statistics public resources (4017 in the 2003-2004 cohort and 3985 in the 2005-2006 cohort).

To calculate a statistical descriptor of each participant’s locomotor activity, we first converted activity counts into discrete states with bin edges *b_k_*, *k* = 1*..K*. Activity level states 1*…K* 1 were then defined as half-open intervals *b_k_ a < b_k_*_+1_, state 0 as *a < b*_1_ and state *K* as *a ≥ b_K_*, where *a* is the activity count value. In this study, we defined 8 activity states with bin edges *b_k_* = *e^k^* − 1*, k* = 1*…*7. Thus, each sample was converted into a track of activity states and a transition matrix (TM) was then calculated for each participant (see below). To ensure that our analysis dealt only with days on which a participant actually performed some physical activity, we applied an additional filter. We excluded days with less than 200 minutes corresponding to activity states *>* 0. Only participants with 4 or more days that passed this additional filter were retained, yielding a total of 11839 samples (age, years: 35 ± 23, range 6 − 84; women: 51%). For PCA and Survival analysis, the only samples used were those for participants aged 40 and older with known follow-up on survival/mortality outcome (age, years: 60 ± 13, range 40 − 84; women: 50%). Once PCA loading vectors were identified, we plotted all NHANES samples’ scores in Fig. 2A, including those for which survival/mortality data were not available.

Transition matrices (TM) *T_ij_, i* = 1*…*8, *j* = 1*…*8 were calculated as a set of transition rates from each state *j* to each other state *i* (the diagonal elements correspond to the probability of remaining in the same activity state). TM elements were calculated as *T_ij_* = *N* (*j*→*i*)*/N* (*j*), where *N* (*j*) is the number of minutes corresponding to state *j* and *N* (*j*→*i*) is the number of times the state *j* was immediately followed by state *i* (in the consecutive minute along the sample record). We next converted the TM from a discrete point map to continuous notation: *W_ij_* = *T_ij_−I_ij_*, where *I_ij_* is the identity matrix. *W_ij_* is the proper TM for which the apparatus of the Markov chains can be used. We used this property to calculate Power Spectral Densities (PSD) and eigenfrequencies (shown in Fig. 1B) based on the assumption that the Markov chain model can be an approximation of observed activity records.

We flattened 8 × 8 TM of each sample into a 64-dimensional descriptor vector for Principal Component Analysis (PCA) and Survival analysis. Additionally, we converted the flattened descriptor to log-scale to ensure approximately normal distribution for elements of the locomotor descriptor (a useful property for the stability of the linear models that we applied in PCA and Survival analysis). All near-zero elements (*<* 10^*−*3^, which corresponds to less than 10 transitions during a week) were imputed by the value of 10^*−*3^ before log-scaling.

### UKB dataset

We accessed data from UK Biobank (UKB) under the approved research project 21988 (formerly 9086). At the time the present study was conducted (2015-2017), locomotor activity data were available for 103710 UKB participants. Physical activity was measured using Axivity AX3 tri-axial accelerometers worn on the wrist for 7 consecutive days. The data were recorded in the low-level format as continuous tracks of 3D acceleration values sampled at 100Hz. Some tracks indicated that hardware errors occurred during the monitoring period. Participants with more than 10 such hardware errors in their track were excluded from our analysis, leaving 102914 participants. To make it possible to apply the PCA and Survival analysis models established using NHANES data to the UKB data, we downsampled the original UKB tracks to 1*min^−^*^1^ (as used in NHANES). For this purpose, individual acceleration records were split into 1-minute slices, and for each slice, the natural logarithm of the sum over the power spectral density (PSD) of the signal within that slice was calculated. Each of these PSDs was calculated from the absolute values of acceleration using the Welch method with 512 points Hann window function and 50% window overlap.

The downsampled UKB tracks represent the level of physical activity per minute but are quantitatively different from the NHANES activity counts. We used a quantile normalization procedure to re-scale the UKB values to the range of discrete activity states of NHANES. We selected NHANES participants in the age range 45-75 and dropped 1.6 of participants with the lowest and highest average activities. The combined tracks from the remaining 2398 NHANES participants were used to calculate the occupancy fractions *p_k_* = *N*(*k*)/*N* for each NHANES activity state (here *N*(*k*) is the number of times the state *k* was seen and *N* is the total number of minutes in all tracks). Then we randomly selected 5000 UKB participants from the same age range and similarly dropped 1/6 of participants with the lowest and highest average activities; this resulted in selection of 3288 UKB participants. Using the combined UKB tracks from selected participants, we found UKB bin edges 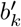 such, that the occupancy fractions for the corresponding activity states, were equal to the occupancy fractions in NHANES. Note that such quantile normalization automatically accounts for shift, linear and monotonic non-linear scaling of values, and so the resulting UKB activity states are roughly equivalent to the ones from NHANES. Once bin edges for UKB were obtained, the downsampled UKB tracks were processed exactly as described above for NHANES. TMs and corresponding descriptors were obtained for 95609 UKB participants (age, years: average 61 *±* 8, median 62, range 43 *−* 78; women: 56%).

### Survival analysis

We estimated hazards ratio using Cox proportional hazards model fit to NHANES 2003 − 2006 linked mortality data. The covariates used in the model included gender label and locomotor activity variables in the form of natural logarithm of transition matrix elements. The total number of covariates was 65 (64 elements of transition matrix and one gender label), so we used regularization parameter *λ* = 0.01. Once fit, the model (“LogMort”) then was applied to produce hazard ratio estimations for NHANES and UKB participants. The model did not explicitly include age of participants. Hazard ratio models for mortality (“LogMort (UKB)”) and morbidity (“LogMorb (UKB)”) were trained in similar way using UKB 3-year follow-up data on death and diagnosis ICD10 codes, respectively. Only UKB participants without any of cardiovascular, diabetes, hypertension, cancer diagnoses at the moment of loocomotor activity measurements were used to train the “LogMorb (UKB)” model (43533 UKB participants with 1331 diseased during 3-year follow-up).

The resulting hazard ratio score was further tested for significance of association with mortality risks again using Cox proportional hazards approach. Now, chronological age and gender were explicitly used as covariates along with the hazard ratio and, optionally, *PC*_1_ score, the latter being an approximation to biological age (see below). Both hazard ratio and *PC*_1_ were linearly detrended by chronological age and gender. This was done to ensure that the obtained significance parameters reflect the contribution of the age-and gender-adjusted part of hazard ratio or *PC*_1_. All procedures were performed in the same way for NHANES and UKB. *PC*_1_ scores for NHANES were obtained using PCA. To obtain *PC*_1_ for UKB participants we calculated projections of UKB variables onto corresponding first eigen vector of NHANES data covariance matrix.

Empirical mortality (i.e. incidence rate depending on age only) was estimated using NHANES death register follow-up data to check consistency with Gompertz law of mortality using parametric Cox-Gompertz proportional hazard model in the form of maximal likelihood optimization adapted from [19] with *M*_0_ and Γ the parameters of Gompertz mortality law, *t_i_*, ∆*t_i_*, and *δ_i_* the age, follow-up time and death event indicator of participant *i*, respectively.

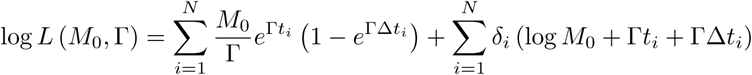

All analyses were conducted using a set of in-house scripts developed in Python [www.python.org] and R [www.r-project.org].

## ACKNOWLEDGEMENTS

This study was conducted using the UK Biobank Resource, application number 21988. We would like to thank G. Ivashkevich, I. Molodtsov, A. Tarkhov, V. Kogan from Gero LLC for extensive help in conducting the research and David K. Edwards for her most valuable help with manuscript editing. Funding: This work was funded by Gero LLC. Author contributions: TP, EG and KA designed and performed the numerical modelling, statistical analysis, wrote the manuscript; BZ collected, analyzed, and interpreted the data; MP performed statistical analysis, wrote the manuscript; LM designed the study, wrote the manuscript; KK and AG wrote the manuscript; PF designed the study, performed the numerical modelling and wrote the manuscript. All authors reviewed the manuscript.

## Appendix A: Transition matrix and Power Spectrum Density

Under the Markov chain model, the evolution of the probability *P_i_* (*t*) to find the system at state *i* for the system with *N* discrete states is governed by the master equation which in the linear mode can be written as

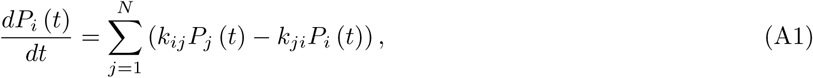

where *k_ij_* ≥ 0 is the rate of transition from state *j* to state *i*. By introducing the transition matrix (TM) according to

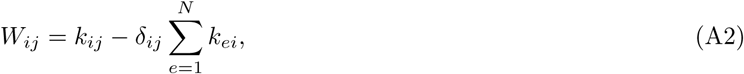

we can rewrite Eq. A1 as

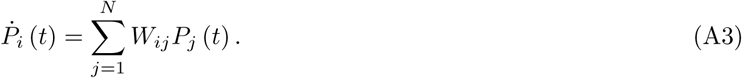

Note from Eq. A2 and definition of *k_ij_* we have *W_ij_ ≥* 0 for *i ≠ j*, *W_ii_ ≤* 0 and

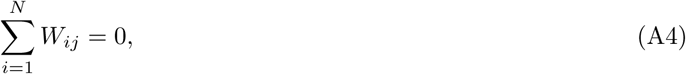

from which it follows that the probability norm is preserved 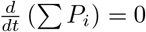, as it should be.

In the following analysis we will assume that the TM *W* is irreducible and has distinct eignevalues. The reasoning for such assumptions will be provided later. Under this assumptions *W* can be diagonalized

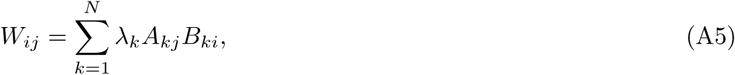

where *A_k_* and *B_k_* are left (*Σ_i_ W_ij_A_ki_*= *λ_k_A_kj_*) and right (*Σ_i_ W_ij_B_ki_*= *λ_k_B_ki_*) eigenvectors corresponding to eigenvalue *λ_k_*. Note that the systems of left and right eigenvectors are the inverse for each other:

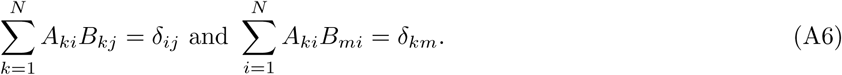

To solve Eq. A3 we introduce

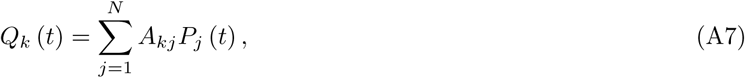

and using Eqs. A5 and A6 rewrite Eq. A3 as

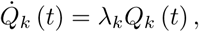

for which the solution is

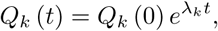

from which using Eqs. A6 and A7 we get

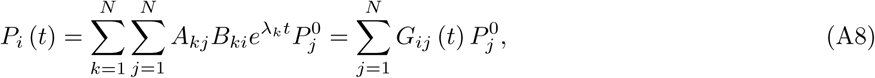

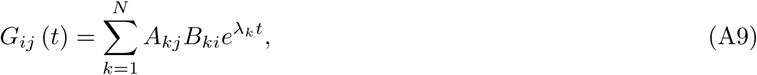

where *G_ij_* (*t*) is the probability *P* (*i, t* |*j,* 0) to find the system at state *i* at time *t* if the system originally was at state *j* at time 0 and *P*^0^ is the initial distribution.

The assumption that *W* has distinct eignavalues together with Eq. A4 imply that *W* has exactly one zero eigenvalue. Since the order of eigenvalues is arbitrary, we can state that

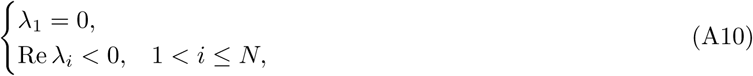

where the later unequality follows from *W* being a TM.

Indeed, *W* is real-valued, therefore for any eigenvalue *λ_i_* and corresponding left eigenvector *A_i_* we have

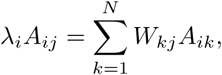

and

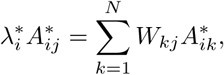

where * is complex conjugate. After multiplying the first equation by 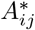, the second by *A_ij_* and summing we get

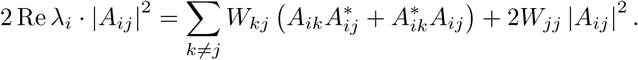

Representing all *A_ij_* in exponential form *A_ij_* = *ρ_ij_* exp (*ιϕ_ij_*), dividing by |*A_ij_* ^2^| and replacing *W_jj_* using Eq. A4 we get

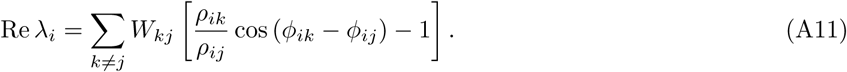

This equation holds for all *i* and *j*. For a given *i* let us choose a particular *j* such that *ρ_ik_* ≤ *ρ_ij_*. Since all *ρ_ik_* and *W_kj_* for *k* ≠ *j* are non-negative by definition, the Eq. A11 becomes Re *λ_i_* ≤ 0.

According to Eq. A3, any equlibrium state is the right eigenvector corresponding to the zero eigenvalue. Since *W* has only one such eigenvector (up to scaling), we have a unique equlibrium distribution given by

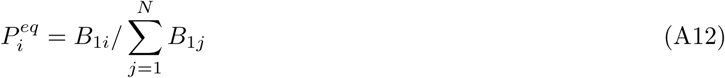

The eigensystem has several interesting properties. From Eqs. A9 and A10 we get *G_ij_* (+*∞*) = *A*_1*j*_*B*_1*i*_ and the distribution at *t* = +*∞* is

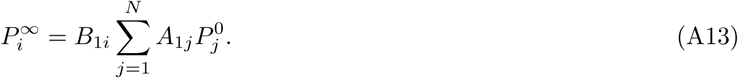

For any initial distribution *P*^0^ the corresponding *P^∞^* is an equlibrium state:

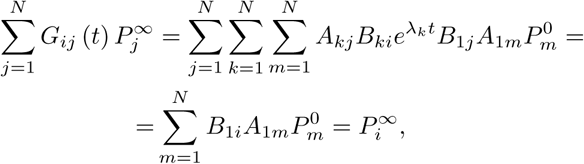

and since equlibrium is unique 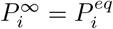 for any *P*^0^. From this and Eq. A13 we have

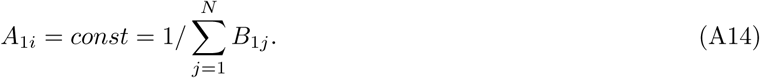

Using Eq. A4, for the right eigenvectors *B_k_* we get

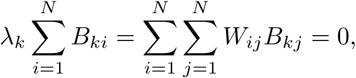

and therefor

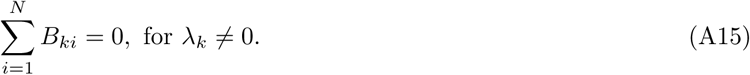

Let as consider a discrete real-valued stochastic process *x* (*t*) having value *x_i_* when system happens to be in state i. According to the Wiener–Khinchin theorem, the power spectral density *S_x_* (*ω*) for the *x* (*t*) is the Fourier transform of the autocorrelator

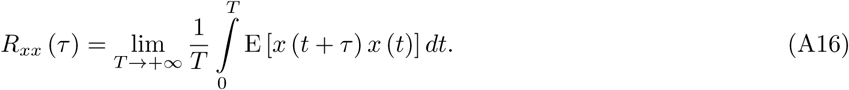

Using the fact that *R_xx_* (*τ*) is an even real-valued function we obtain

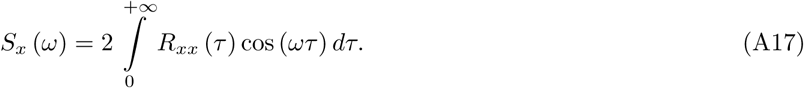

Here we follow the common physical convention that the total power of the signal is given by 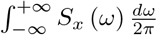.

Expanding the Eq. A16 we get
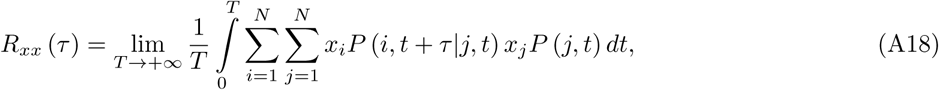

where *P* (*i, t* + *τ |j, t*) is the probaility to find the system at state *j* at time *t* + *τ* if the system was at state *j* at time *t* and *P* (*j, t*) is the probability to find the system at state *j* at time *t*, with the evolution of the system starting from some state *P*^0^. From the definitions we have *P* (*i, t* + *τ |j, t*) = *G_ij_* (*τ*) for *τ* ≥ 0 and *P* (*j, t*) = *P_j_* (*t*). Using this and Eqs. A8 and A9 rewrite Eq.A18 as

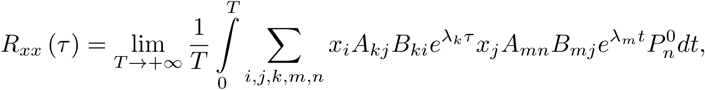

where *τ >* 0 and the summation is done for each index from 1 to *N*. By rearranging and using Eq. A10 we get

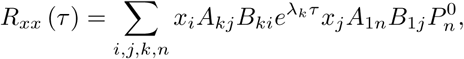

from which using Eq. A12 we finally obtain

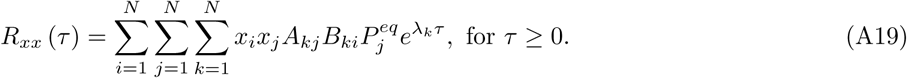

Note that *R_xx_* (*τ*) is not dependent on the initial distribution *P*^0^, as it is expected for the system with equlibrium state. The integration of Eq. A19 using Eq. A17 is straightforward, and we get

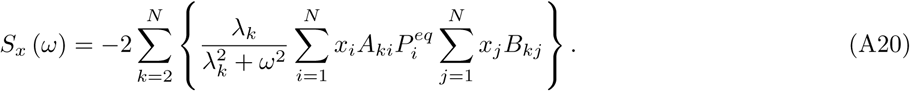

The Eq. A20 is valid for any irreducible diagonalizable TM *W*. In particular, some *λ_k_* may be complex. However, for the real-valued matrix *W* complex eigenvalues and corresponding eigenvectors always comes in complex conjugate pairs, which, together with Eq.A10, imply that *S_x_* (*ω*) is always real positive, as any PSD should be.

Due to time symmentry of the fundamental physical laws, for the systems in thermodynamic equilibrium the detailed balance assumption is hold:

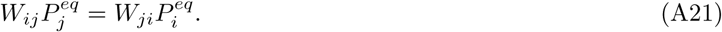

Biological organisms as a whole are not systems in thermodynamic equlibrium and the description of the motion using Markov chain model is only a rough approximation, so there are no *a priori* reasons to assume the detailed balance. However, experimetally the correlation between 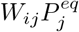 and 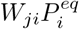 is good, so it is interesting to see how *S_x_* (*ω*) looks under detailed balance assumption.

First we introduce a derived matrix

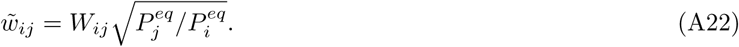

With Eq. A21 hold, *w̃* is symmetric and therefore can be eigendecomposed into

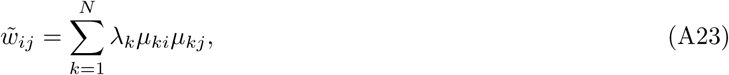

where all eignvalues *λ_k_* are real and eigenvectors *μ_k_* are orthonormal:

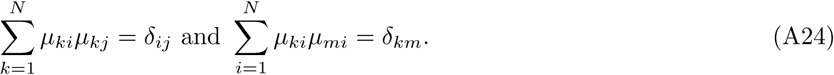

From Eqs. A22 and A23 we get

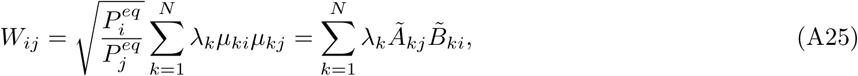

where

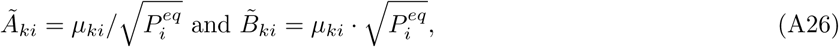

from which using Eq. A24 follows

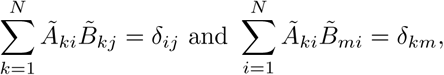

which imply that Eq. A25 is an eigendecomposition of *W* as in Eq. A5, so we can use Eq. A20, which becomes

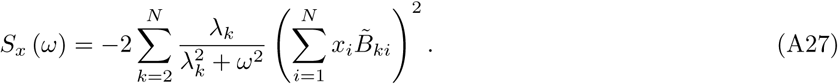

Here we used 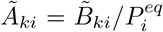, obtained from Eq. A26, to express *S_x_* (*ω*) via right eigenventors *B*̃*_k_* alone. Each of the right eigenvectors *B_k_* is defined up to a multiplication constant, however the scaling is fixed for *B*̃_*k*_: from Eqs. A24 and A26 we have

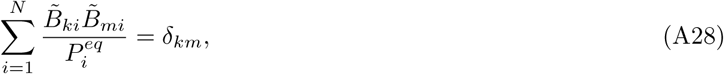

from which we can find a proper scaling for an arbitrary right eigenvector *B_k_*:

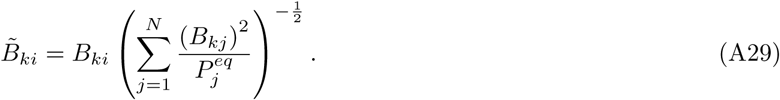

The *S_x_* (*ω*) can be calculated under detailed balance assumption as follows: calculate the right eigensystem for *W*, scale the found eigenvectors *B_k_* using Eq. A29 and finally calculate *S_x_* (*ω*) using Eq. A27. The same procedure can be applied when the detailed balance assumption holds only approximately, as long as we drop the imaginary part of the found eigenvalues and right eigenvectors. Note that even when all eigenvalues are real, the Eq. A27 is not equivalent to Eq. A20 without the detailed balance assumption. In particular, scaling according to Eq. A29 is not enought for Eq. A28 to hold, which is required for Eq. A27 to be precise.

